# Alternative strategy to induce CRISPR-mediated genetic changes in hematopoietic cells

**DOI:** 10.1101/2021.10.25.465699

**Authors:** E González-Romero, A Rosal-Vela, A Liquori, C Martínez-Valiente, G García-García, JM Millán, MA Sanz, JV Cervera, RP Vázquez-Manrique

**Author notes:** **Correspondence:** José Vicente Cervera, PhD, MD, Hematology Department/Genetics Unit, IIS-Hospital La Fe, Avda. de Fernando Abril Martorell, n° 106. 46026 Valencia, Spain, Rafael Vázquez-Manrique, PhD, Molecular, Cellular and Genomic Biomedicine, IIS-Hospital, La Fe, Avda. de Fernando Abril Martorell, n° 106. 46026 Valencia, Spain, /1246685. Authors contributed equally (JV Cervera, RP Vázquez-Manrique).

## Abstract

Acute Myeloid Leukaemia is a complex heterogenous disease caused by clonal expansion of undifferentiated myeloid precursors. Recently, several haematological models have been developed with CRISPR/Cas9, using viral vectors, because blood cells are hard to transfect. To avoid virus disadvantages, we have developed a strategy to generate CRISPR constructs, by means of PCR, which any lab equipped with basic technology can implement. These PCR-generated constructs enter easily into hard-to-transfect cells. After testing its functionality by editing *MYBL2* gene in HEK293 cells, we successfully introduced the R172 mutation in *IDH2* gene in NB4 cells that expresses constitutively the Cas9 nuclease. Comparing our methodology with ribonucleoprotein strategies, we found that mutation introduction efficiency was similar between both methodologies, and no off-target events were detected. Our strategy represents a valid alternative to introduce desired mutations in hard to transfect leukemic cells, avoiding using huge vectors or viral transduction.

## INTRODUCTION

CRISPR are repetitive DNA sequences that, together with the CRISPR associated proteins (Cas proteins), act as a prokaryotic adaptive natural defence against virus attack described in archaea and eubacteria [1–4]. The main components of this system have been adapted to be used ectopically to edit all sorts of cells and organisms [5–9]. As a consequence, the CRISPR technology has emerged as a revolutionary way to manipulate genomes of all types, through animal models and plants, to human cells. Using CRISPR make it possible to develop several haematopoietic cell models (reviewed by González-Romero *et al*. [10]). However, the main handicap of these cells is the poor transfection efficiency. Thus, viruses are preferred to introduce CRISPR into cells.[11–14] These methods present disadvantages ^like^ packing capacity, induction of immune reactions or integration of lentivirus [15].

Acute myeloid leukaemia (AML) is a heterogeneous disease produced by clonal expansion of myeloid precursors, resulting in impaired haematopoiesis and bone marrow failure [16]. Mutations in the isocitrate dehydrogenase 2 enzyme gene (*IDH2)* are associated with specific clinical outcomes and characteristic gene expression and epigenetic changes [17]. *IDH2* presents two recurrent mutations, *IDH2*^*R140*^ and *IDH2*^*R172*^. *IDH2*^*R172*^ has been proposed as a trait to create a new category of AML [18]. Therefore, studying these mutations using cell and animal models is essential to provide with tools to find out new therapies against AML. In this regard, CRISPR bring us an invaluable toolkit to create these models.

In this study, we developed a strategy to easily generate constructs to introduce CRISPR/Cas9 elements, in an efficient way, into the difficult-to-transfect NB4 haematological cells. We designed an easy way to assemble small constructs to express single RNA guides (sgRNAs). Alternatively, these constructs include GFP, to follow transfection or/and to isolate singled cells. We generated a NB4 cell line that constitutively expresses Cas9 (NB4-Cas9), that allowed us to introduce our easy-to-make constructs to produce the *IDH2*^R172^ mutation. Moreover, we proved the effectiveness of our strategy successfully cleaving another gene target, *MYBL2*. This gene encodes a transcription factor involved in cell cycle, cell survival and cell differentiation regulation [19]. Deregulations in its expression are related with a broad spectrum of cancer entities, as AML [20]. Finally, we compared the efficiency of our new strategy with ribonucleoprotein complexes, and we did deep sequencing analysis to analyze the efficiency and to discard off-target events.

## MATERIAL AND METHODS

### Construction of the pEGR1 vector

To clone the sgRNA cassette (pU6 promoter, sgRNA scaffold and terminator) in pEGFP-N1, primers with AflII restriction sites were used to amplify sgRNA cassette from the PX458 plasmid (Addgene 48138) [21]. pEGFP-N1 and insert were digested with AflII (Thermo Fisher Scientific, Waltham, Massachusetts, US). Digested vector was dephosphorylated with Alkaline Phosphatase (New England Biolab, Ipswich, Massachusetts, United States). Then, both were purified using QUIAquick PCR Purification Kit (QIAGEN, Hilden, Germany) and ligated by T4 DNA ligase (Thermo Fisher Scientific) at RT. The product was electroporated in Top10 electrocompetent cells. We used PCR to screen for positives. The plasmid from these bacterial clones was purified with QIAprep Spin Miniprep Kit (QIAGEN) and verified by sequencing (Figure 1A).

**Figure 1.**
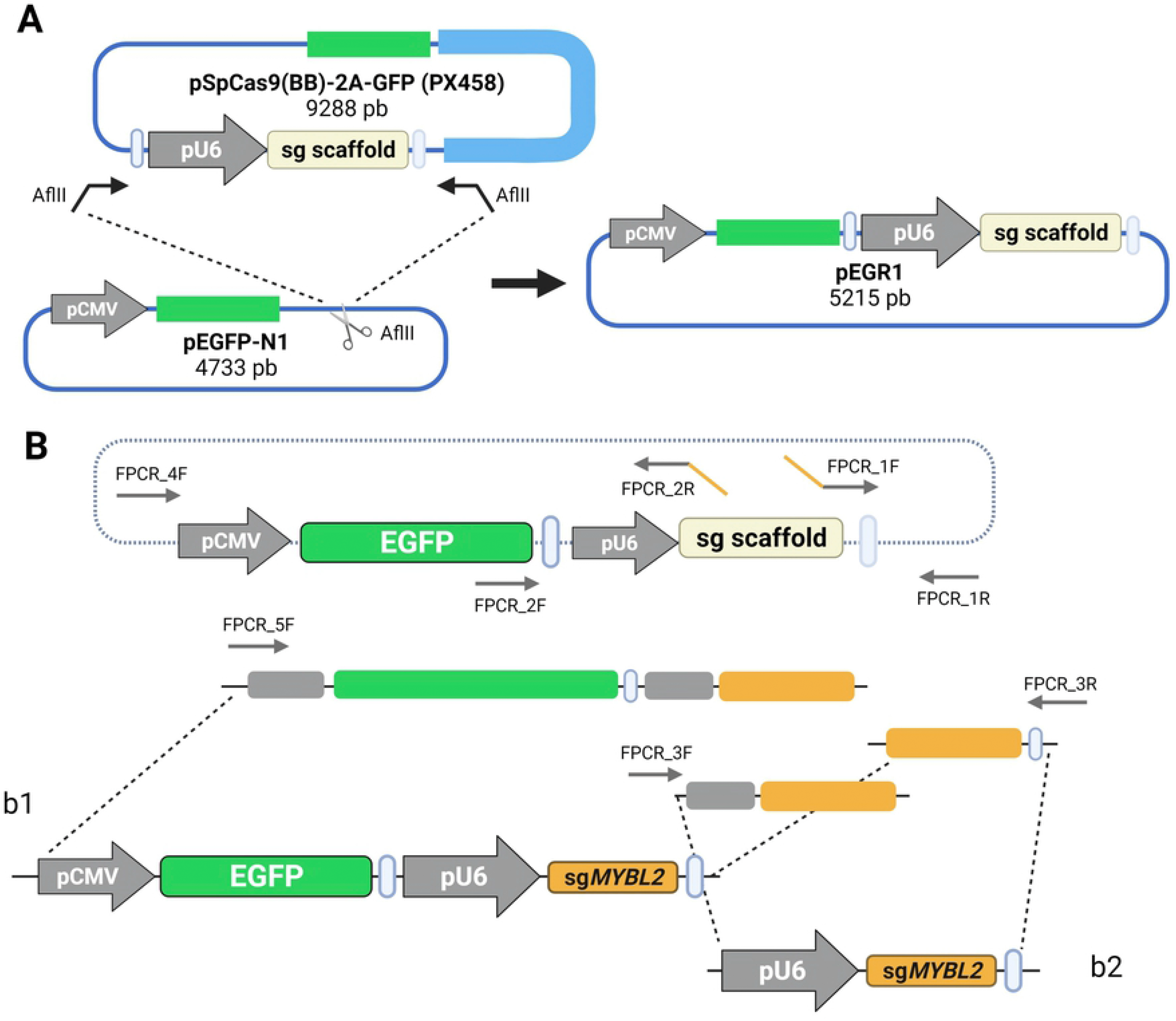
PCR-generated sgRNA constructs. **(A)** Development of the pEGR1 vector, as a template to generate sgRNA constructs by PCR. The sgRNA cassette was amplified from the pX458 vector and then was inserted in the pEGFP-N1 vector. **(B)** Schematic illustration of the pEGFP-pU6-sgRNA scaffold plasmid, pEGR-1. With specific primers it is possible to generate a construct, using fusion PCR, containing either the GFP cassette and the sgRNA (b1), or just the sgRNA (b2). First, to create the pU6-sgRNA module, the PCR_1F and PCR_1R primers were used to amplify the terminator sequence with a specific sgRNA. FPCR_1F presents in 5′ the specific 20 nucleotide sequence (orange). The region of the vector in which the FPCR_1R primer lies is complex and contains repetitions. To avoid this issue, we inserted a tail in 5′ of this primer, that will be used as a template in subsequent PCR reactions. FPCR_2F and FPCR_2R primers amplify U6 promoter and a specific sgRNA. In that case, a part of the sequence of FPCR_2R is the sgRNA reverse sequence. These two products were merged in a specific PCR with FPCR_3F and FPCR_3R primers because both products overlap (b1). Following the same strategy, the EGFP-U6-sgRNA module was created using FPCR_4F and FPCR_4R primers to amplify CMV promoter, *EGFP* gene, terminator and the sgRNA reverse sequence. At the same way that previous construct, these two constructs can be merged. In this case FusionPCR_5F and FusionPCR_3R primers are used. FusionPCR_3R has homology against FusionPCR_1R tail, so it is used in both fusions (b2).

### Creating the sgRNA constructs by fusion PCR

Different primer combinations were used to create the constructs that encode the sgRNAs against *IDH2* or *MYBL2*. All PCRs were carried out using the Phusion High-Fidelity Polymerase (Thermo Fisher Scientific) and pEGR1 as a template. Detailed protocol to create pU6-sgRNA and EGFP-U6-sgRNA modules is explained in Figure 1B. We phosphorylated the primers with T4 Polynucleotide kinase to generate the phosphorylated sg*MYBL2* construct (New England Biolab). After phosphorylation, primers were purified by MinElute PCR Purification Kit (QIAGEN), to do the fusion PCRs. All constructs were also purified by MinElute PCR Purification Kit (QIAGEN) prior transfection.

### Design of sgRNAs and ssODN

The sequence around the target in *IDH2* was sequenced from NB4 cells by Sanger Sequencing to ensure that there was no single nucleotide polymorphism (SNPs) present, that may prevent from homologous recombination events. This sequence was used as a bait in sgRNA predictor web page chopchop.cbu.uib.no [22]. To test the *MYBL2* gene, we designed a sgRNA in the intron 3 of the gene, used before in the lab. To introduce the R172K mutation, a single stranded DNA (ssODN) was designed with 35 nt homology arms, the *IDH2*^R172K^ point mutation and silent changes to prevent Cas9 reiterated cleavage. The sg*IDH2*_2 PAM silent change introduces a new point of cleavage site for the restriction enzyme HhaI (New England BioLabs). For the sg*IDH2*_1 PAM it was impossible to modify the essential sequence, so we introduced seven silent changes in sg*IDH2*_1 sequence (Figure 3). The ssODN was synthesized as Ultramer Oligonucleotides (IDTDNA, Coralville, Iowa, United States).

**Figure 2.**
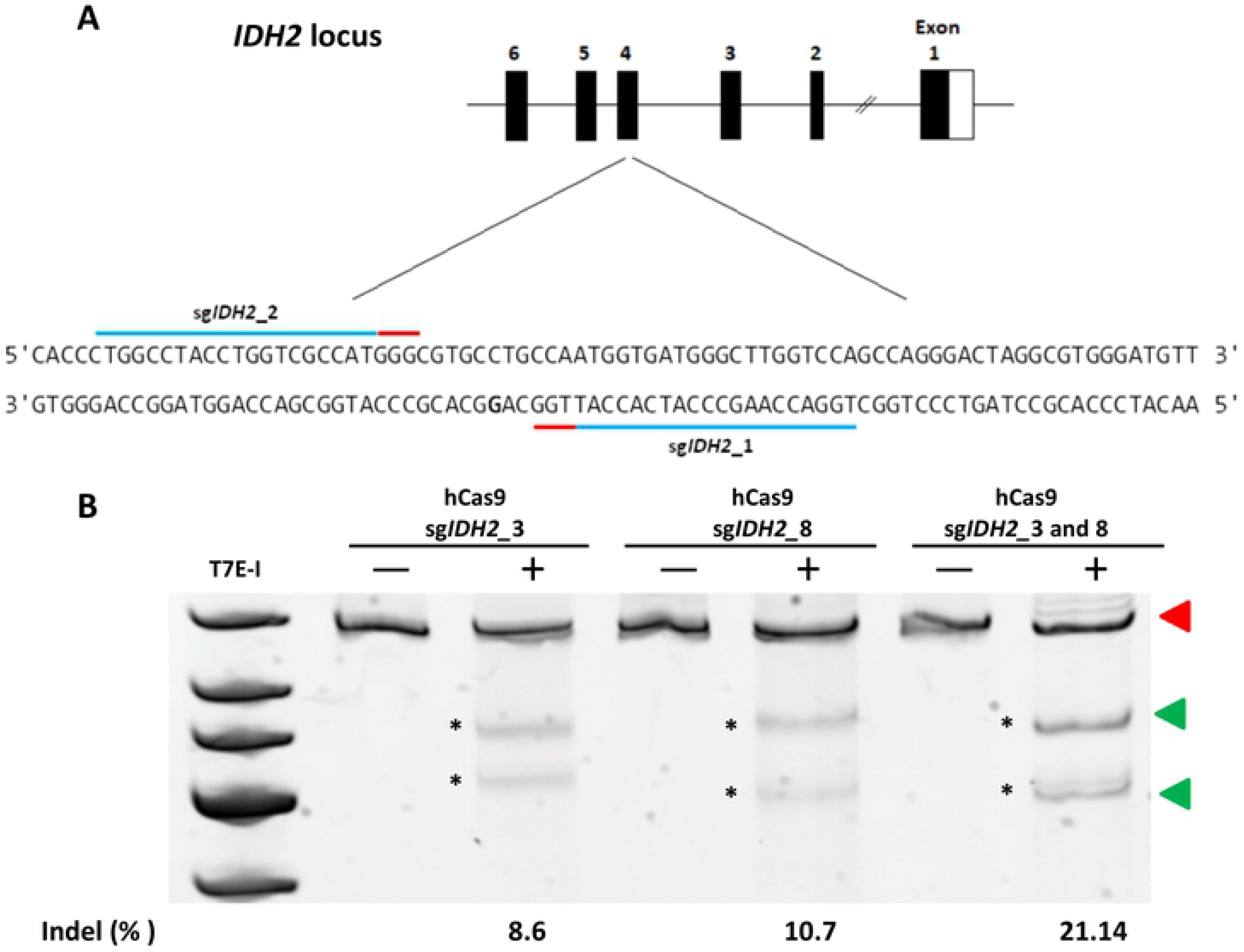
Targeting *IDH2* gene in HEK293 cells. **(A)** Structure of *IDH2* gen, in detail the position of R172K mutation in exon 4. Blue lines indicate designed sgRNA, and red lines indicate the protospacer adjacent motif (PAM) sequences. (**B)** T7 endonuclease I assay depicting indel formation by candidate sgRNAs. Black numbers show band relative intensities (%). Red triangles point uncuted products, and green triangles T7E-I products.

**Figure 3.**
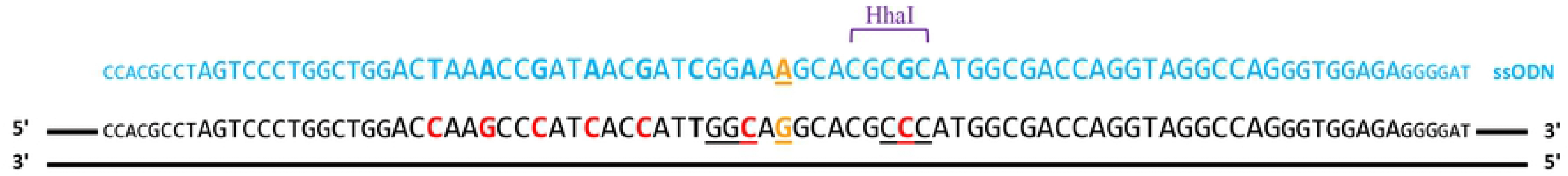
Diagram of the ssODN designed to target *IDH2*. The R172 mutation is indicated in orange and additional changes were introduced to avoid Cas9 re-cutting (red). The change introduced in the PAM sequence (underlined nucleotides) of the sgRNA_2 introduce the HhaI target sequence, that is used for the RFLP analysis of the editing efficiency.

### PCR amplified Cas9 cassette

The whole hCas9 cassette PCR from hCas9 vector was amplified by Phusion High-Fidelity DNA Polimerase (Thermo Fisher Scientific) with specific primers. Before cell transfections, PCR Cas9 was purified by MinElute PCR Purification Kit.

### PX458-sgMYBL2 vector

PX458 with sg*MYBL2* expression was produced following a previous protocol [21].

### HEK293 cell culture and transfections

HEK293T and HEK293 cells were cultured in DMEM 1X (Thermo Fisher Scientific) supplemented with 0.5 % Peniciline/Streptomycin 100X Solution (Biowest, Minneapolis, U.S.A) and 10% Fetal Bovine Serum (Thermo Fisher Scientific). Cells were incubated at 37°C at 5 % CO_2_. For CRISPR transfections we used Lipofectamine3000 (Invitrogen), 0.9 x10^5^ cells and pMAX was transfected as control. For sg*MYBL2* and hCas9 plasmid (Addgene 41815) [23] co-transfection, we tested different concentrations: 500 ng hCas9– 23.3 ng sg*MYBL2*; 250 ng hCas9-12 ng sg*MYBL2*; 250 ng hCas9-35 ng sg*MYBL2*; 250 ng hCas9-60 ng sg*MYBL2* and 150 ng hCas9-7 ng sg*MYBL2*. To test the effect of the 5′ termini phosphorylation, phosphorylated sg*MYBL2* construct was transfected together with the hCas9 vector. Finally, we assayed transfection of 35 ng of sg*MYBL2* PCR construct together with 250 ng of the PCR-amplified Cas9 cassette.

For *IDH2* editing, 35 ng of each guide or 17.5 ng of both guides were co transfected with 250 ng of hCas9 vector. For editing experiments, 10 μM of ssODN was used.

### Generation of the NB4 cells expressing constitutively Cas9

One day before transfection, 3×10^6^ 293T cells were seed in a 10 cm dish. Cells were transfected with lentiCRISPR V.2 and two packaging plasmids (pPAX2 and pMD2.G) with the Cl_2_Ca method. 48 h post transfection, the supernatant containing the lentivirus was collected, filtered through a 0.45 μm nitrocellulose filter and stored at −80°C for further use. The lentiCRISPR V.2 contains the *Streptococcus pyogenes* Cas9 nuclease and puromycin resistance gene. For lentiviral transduction, 5 x10^5^ NB4 cells were infected with 100 μL of the lentivirus supernatant and polybrene to a final concentration of 4 μg/μL. After 24 h, cells were centrifuged and resuspended in media with puromycin (0.6 μg/mL) (InvivoGen, San Diego, California, U.S.A). Cells were maintained with these conditions several days. To isolate positive cells (i.e. with inserted virus) DNA was extracted to detect the presence of the Cas9 gene by PCR. After PCR confirmation, NB4 Cas9 positive were maintained in media supplemented with 0.2 μg/mL of puromycin.

### NB4 cell culture and nucleofection

NB4 cells were cultured in RPMI Medium 1640 (Gibco, Thermo Fisher Scientific), supplemented with 1% Peniciline/Streptomycin 100X Solution and 10% Fetal Bovine Serum. NB4-Cas9 medium is supplemented with 0.2 ug/mL of puromycin. Cells were incubated at 37 °C and with 5 % CO_2_. For nucleofection experiments we used Cell Line Nucleofector Solution Kit V (Lonza Basel, Switzerland), 2×10^5^ cells and pMAX was transfected as a positive control. We followed Lonza Protocol for Kit V. NB4-Cas9 cells were nucleofected with 400 ng or 800 ng of sg*IDH2*_1 and sg*IDH2*_2 constructs. To assemble the ribonucloprotein (RNP)-ensembled complexes, the crRNA and tracrRNA (IDTDNA) elements were hybridized to form sgRNA following manufacturer`s protocol. Prior to nucleofection, 17 μg of purified recombinant *S. pyogenes* Cas9 nuclease (IDTDNA) was added to 20.3 μg of each duplex and incubated for 10 min at RT to form RNP complex. Two hundred thousand NB4 cells were nucleofected. For edition experiments 1uL of ssODN (100 μM) was used.

### On-target and off-target analysis by Next Generation Sequencing

Using Cas-OFFinder [24] we selected potential off-targets that differed from sg*IDH2*_1 and sg*IDH2*_2 by up tree mismatches (Table 1). The same samples used for RFLP assays were used as templates to analyse on-target and the 16-potential off-targets. Two step PCR strategy was used to generate the library. For the first PCR, primers for each locus contained an adapter sequence. PCR products were purified with AMPure Beads (BD Bioscience, Franklin Lakes, New Jersey, US). For the second, PCRs were re-amplified with primers containing the adapter sequence overlapping the first primers, and with an index sequence in the reverse primers. Final PCR products were purified with AMPure Beads (BD Bioscience). The library was prepared with PCR products pooled in equimolar amounts following the manufacture’s protocol and loaded in a Micro MiSeq Reagent Kit v2 (500-cycles) (Illumina, San Diego, CA, USA) on a MiSeq platform (Illumina). The fastaq.gz files were analysed through CRISPResso2 software [25] to evaluate the editing efficiency and possible off target effects, using default parameters.

**Table 1.**
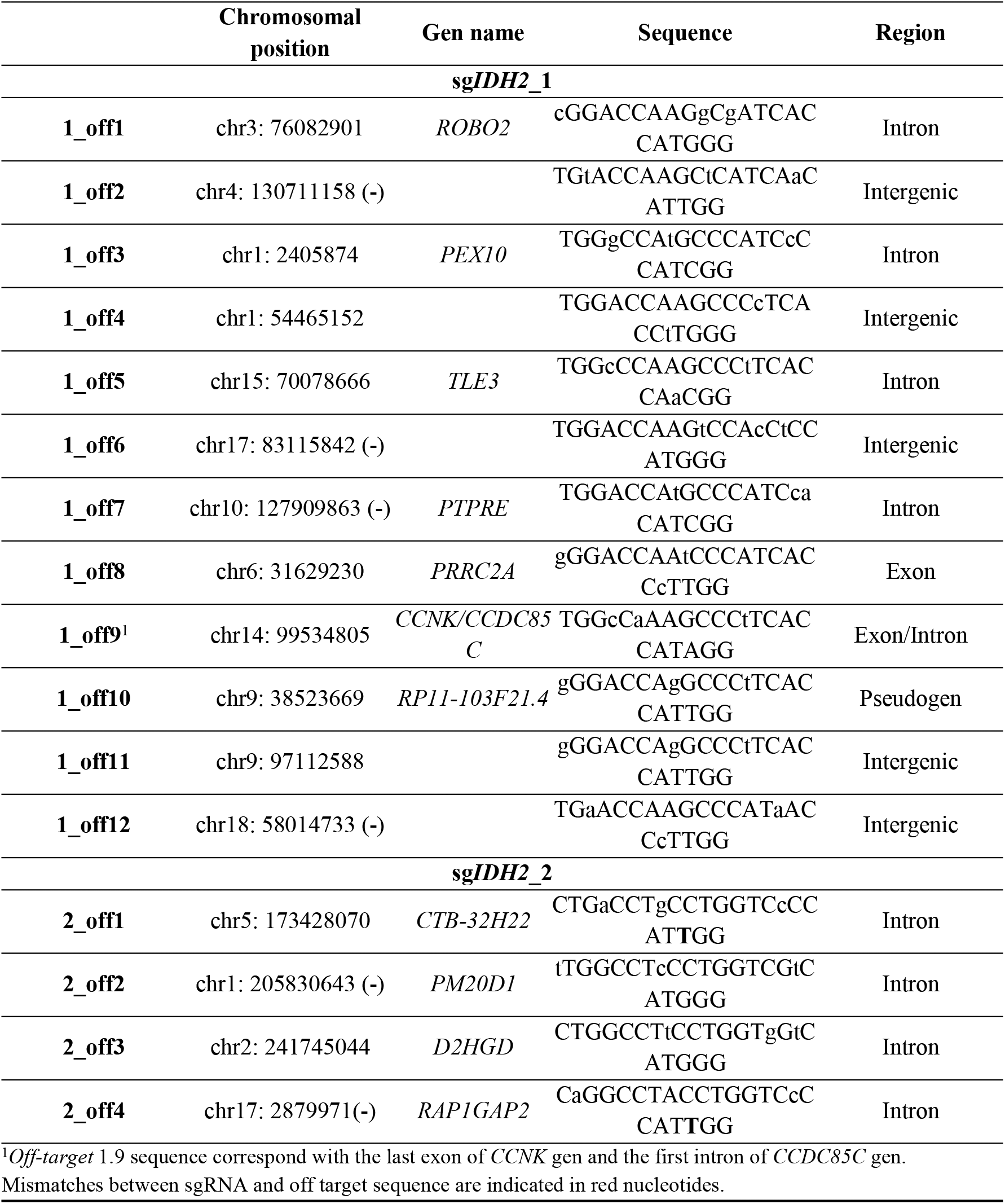
Off-targets selected.

### FACS

Rates of GFP+ transfected cells with pMAX, pX458 vector and GFP-sgMBYBL2 construct were measured by flow cytometry. We nucleofected 800 ng of pX458, 800 ng of GFP construct and 500 ng of pMAX in 2 x10^5^ NB4-Cas9 cells. Cells were analysed on a FACSDiva 8.0.1 (BD Bioscience) 24 h after nucleofection.

### T7 Endonuclease I and RFLP analysis

To evaluate Cas9 cleavage and editing efficiency with T7-Endonuclease I (T7-EI) or RFLP analysis, the genomic DNA of transfected cells was extracted 48 h post transfection. For both protocols, target cleavage sites were amplified by PCR with specific primers. After agarose purification with E.Z.N.A Gel Extraction Kit (OMEGA, Biel/Bienne, Switzerland), T7-EI (New England Biolabs) assays were carried as is described in literature [26]. For RFPL assay, HhaI manufacture’s protocol was carried. Products of both protocols were separated by DNA 10% polyacrylamide or 2% agarose gels and stained with SYBR Safe DNA Gel Stain (Thermo Fisher Scientific). ImageJ program was used to estimate indel or editing percentage measuring integrated intensity of digested and undigested products.

## RESULTS

### PCR-generated sgRNAs are functional in HEK293 cells

To develop an efficient, affordable and general strategy to edit leukemic cell lines we sought to create DNA constructs to deliver sgRNAs as small as possible with better chances to get into cells. Firstly, we created pEGR1, containing structural sequences to induce expression of the sgRNAs (i.e. promoter, the conserved sequence of the sgRNA and poly (A)-signal) but lacking the specific target sequence. From pEGR1 it is possible to produce two kind of constructs by PCR. One carrying the sgRNA for the specific target (465 bp), and another with the guide and EGFP (2033 bp). In this way, we generated sgRNAs with *MYBL2* guide and with *IDH2*_1 and *IDH2*_2 guides.

Before testing our system in leukemic cells, we checked the functionality of the PCR-engineered sgRNA constructs in HEK293 cells. First, we explored the cut efficiency co-transfecting sg*MYBL2* constructs with the hCas9 vector. Different construct/vector ratios were assayed to optimize the experiments. These experiments showed that increasing the amount of DNA produced a dose-dependent increase in non-homologous end joining (NHEJ) events. However, from 250 ng of hCas9 and 35 ng of sgRNA, there was no further increase (10.3 %), despite we increased the quantity of hCas9 and sg*MYBL2*. So, we selected this mix of components for further experiments. Comparing these results with the transfection of the PX458 vector with the same *MYBL2* guide, we found that the cut efficiency obtained with this vector (7.8 %) is lower to the cut efficiency produced with sg*MYBL2* construct (Table 2).

**Table 2.** Results of indels obtained in the different optimizations probed with sg*MYBL2* in HEK293 cells.

| Conditions | 150 ng<br>hCas9<br>7 ng<br>sgMYBL2 | 250 ng<br>hCas9<br>12 ng<br>sgMYBL2 | 250 ng<br>hCas9<br>35 ng<br>sgMYBL2 | 250 ng<br>hCas9<br>60 ng<br>sgMYBL2 | 500 ng<br>hCas9<br>23.3 ng<br>sgMYBL2 | 250 ng<br>hCas9<br>35 ng<br>sgMYBL2<br>-P | 250 ng<br>PCR<br>Cas9<br>35 ng<br>sgMYBL2 | PX458-MYBL2 |
| --- | --- | --- | --- | --- | --- | --- | --- | --- |
| Exp.1 | 4.5 | 3.45 | 5.2 | 6.0 | 5.2 | 14.0 | 3.0 | 4.3 |
| Exp. 2 | 3.0 | 5.8 | 8.0 | 6.0 | 3.0 | 8 | 7.0 | 11.0 |
| Exp. 3 | 3.7 | 4.0 | 13.0 | 14.7 | 14 | 6 | 2.4 | 3.0 |
| Exp. 4 | 2.5 | 4.0 | 14.8 | 5.3 | 8.4 | 13.9 | - | 13.0 |
| Average Indel<br>(%±SEM) | 3.4±0.4 | 4.3±0.5 | 10.3±2.2 | 8.0±2.2 | 7.6±2.3 | 10.5±2.0 | 4.1±1.4 | 7.8±2.5 |

Then, we reasoned that a smaller DNA construct encoding Cas9, rather than the average CRISPR plasmid encoding the plasmid replication components, would enter easier into cells. Hence, we amplified the whole Cas9 cassette (i.e. CMV promoter, Cas9 gene, and polyA-signal) from the hCas9 plasmid. This PCR product is around half the size (≈ 5000 bp) of the plasmid (9553 bp). The PCR products of Cas9 were functional in HEK293 cells but the cut efficiency was substantially reduced compared to the experiments described above (4.1 %) (Table 2).

Previously, other researchers suggested that PCR products with phosphorylated 5′ ends induce enhanced expression of their encoded products [27]. Thus, we transfected sg*MYBL2* with 5′ phosphorylated ends, to study the effect in targeting efficiency, but no differences were found with non-phosphorylated constructs (Table 2). Considering previous optimizations, we continued exploring construct functionality on the *IDH2* gene. sg*IDH2*_1, sg*IDH2*_2 (Figure 2A) and both constructs together were transfected with the hCas9 vector following the selected ratio above. Co-transfection of both sgRNA showed the highest indel production (21.1 %), compared to individual sgRNAs (10.8 % for sg*IDH2*_1 and 8.3 % for sg*IDH2*_2) (Figure 2B) (Table 3). With both sgRNAs, we tried co-transfection with the PCR-amplified Cas9 cassette, but we found a substantial indel reduction (3.36 %) (Table 3).

**Table 3.** Indels and editing results obtained for *IDH2* edition optimization in HEK293 cells.

| Conditions | sgIDH2_1 | sgIDH2_2 | sgIDH2_1 +<br>sgIDH2_2 | 250 ng PCR<br>Cas9<br>sgIDH2_1 +<br>sgIDH2_2 | sgIDH2_1 +<br>sgIDH2_2<br>10 μM ssODN |
| --- | --- | --- | --- | --- | --- |
| Exp.1 | 19.0 | 7.1 | 24.2 | 4.45 | 0.5 |
| Exp. 2 | 11.0 | 15.0 | 28.8 | 2.82 | 0.865 |
| **Exp. 3** | 9.0 | 5.4 | 11.8 | 2.8 | 1.165 |
| **Exp. 4** | 4.0 | 5.5 | 19.7 | - | - |
| **Average indel**<br>**or edition**<br>**(%±SEM)** | 10.8±3.1 | 8.3±2.2 | 21.1±3.6 | 3.36±0.55 | 1.01±0.19 |
| For each experiment, 250 ng of hCas9 vector and 35 ng of guide construct were used, less in PCR Cas9 experiments. |  |  |  |  |  |

Finally, to introduce the *IDH2*^R172^ mutation we used ssODN as a template carrying the point mutation, six silent modifications to avoid further events of DNA cleavage and a restriction site for HhaI (Figure 3). The ssODN was co-transfected at 10 μM with *IDH2* guides and hCas9 vector. By RFLP assay with HhAI enzyme we got an editing efficiency of 1.01 % (Table 3).

### PCR products encoding sgRNAs are functional and readily transfected into leukaemia cell lines

To compare the efficiency of our sgRNA with the available CRISPR plasmids, we nucleofected the EGFP-sg*MYBL2* constructs, the small commercial vector pMAX and the PX458 plasmid in the NB4-Cas9 line. Then, the EGFP positive cells were measured by flow cytometry. As we expected, while PX458 plasmid produced only 1 % of EGFP positive cells, EGFP-sg*MYNL2* produced almost 40 %, while with pMAX vector we obtained almost 26 % (Figure 4). We observed no significant cell death, hence we believe that there is not toxicity from nucleofection experiments.

**Figure 4.**
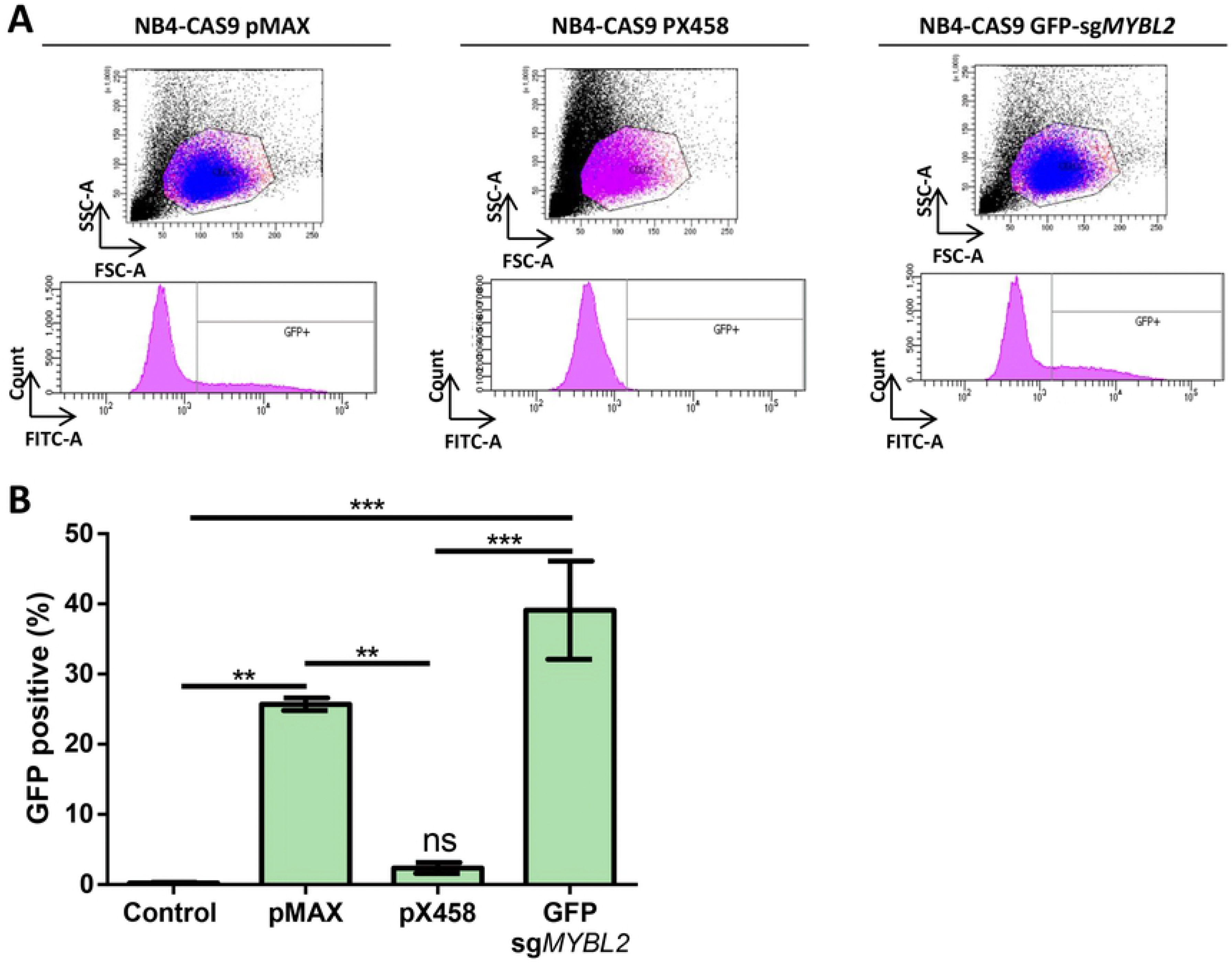
PCR-generated sgRNAs gets easily into NB4 cells. **(A)** Cytometry analysis of NB4 cells transfected with GFP-sgMYBL2 construct, shows that they present a similar number of GFP positive cells (blue color) than cells transfected with the commercial pMAX vector. In contrast, we observe very little transfection efficiency using the PX458 plasmid. **(B)** Using the EGFP-sgMYBL2 PCR product we obtained 39.1%±7.0 (Mean±SEM) of EGFP+ cells). This percentage was higher than the EGFP+ cells produced with transfection of the small pMAX vector, 25.7%±0.9, although the difference was not statistically significant (one-way ANOVA test, with posthoc Tukey). Transfection with these constructs were statistically different from the fluorescent background (0.2%±0.1) with p-values <0.01 and 0.001 (pMax and EGFP-sgMYBL2, respectively) Both, pMAX and EGFP-sgMYBL2 showed statistically differences with the transfection results obtained with pX458 (2.4%±0.8), with p-values <0.01 and 0.001 respectively.

Then, we focused on the *IDH2* gene to optimize targeting and edition in NB4-Cas9 cells. First, we transfected the *IDH2* constructs 1 and 2 together, at two different equimolar concentrations (800 and 1500 ng). Although the differences were not statistically significant, we observed a slightly higher efficiency of indel production with the highest concentration (14.52 %) (Figure 5A). To introduce *IDH2*^R172^ we optimized the ssODN concentration, which resulted to be 100 μM, which produced 2.2 % of edition (Table 4) (Figure 5B).

**Table 4.** Indel and editing efficiencies using sg*IDH2* constructs in NB4-Cas9 cells and RNPs in NB4 cells.

| Conditions | sgIDH2_1 +<br>sgIDH2_2 | RNPs | sgIDH2_1 +<br>sgIDH2_2<br>100 μM | RNPs<br>100 μM |
| --- | --- | --- | --- | --- |
| Exp. 1 | 21.56 | 28.7 | 2.8 | 2.5 |
| Exp. 2 | 7 | 34.16 | 2.3 | 1.6 |
| Exp. 3 | 15 | 26.7 | 1.5 | 2.0 |
| Average indel<br>or edition<br>(%±SEM) | 14.52±7.28 | 29.85±3.86 | 2.2±0.4 | 2.0±0.3 |
| For editing NB4-Cas9 cells, 750 ng of each constructs were used. |  |  |  |  |

**Figure 5.**
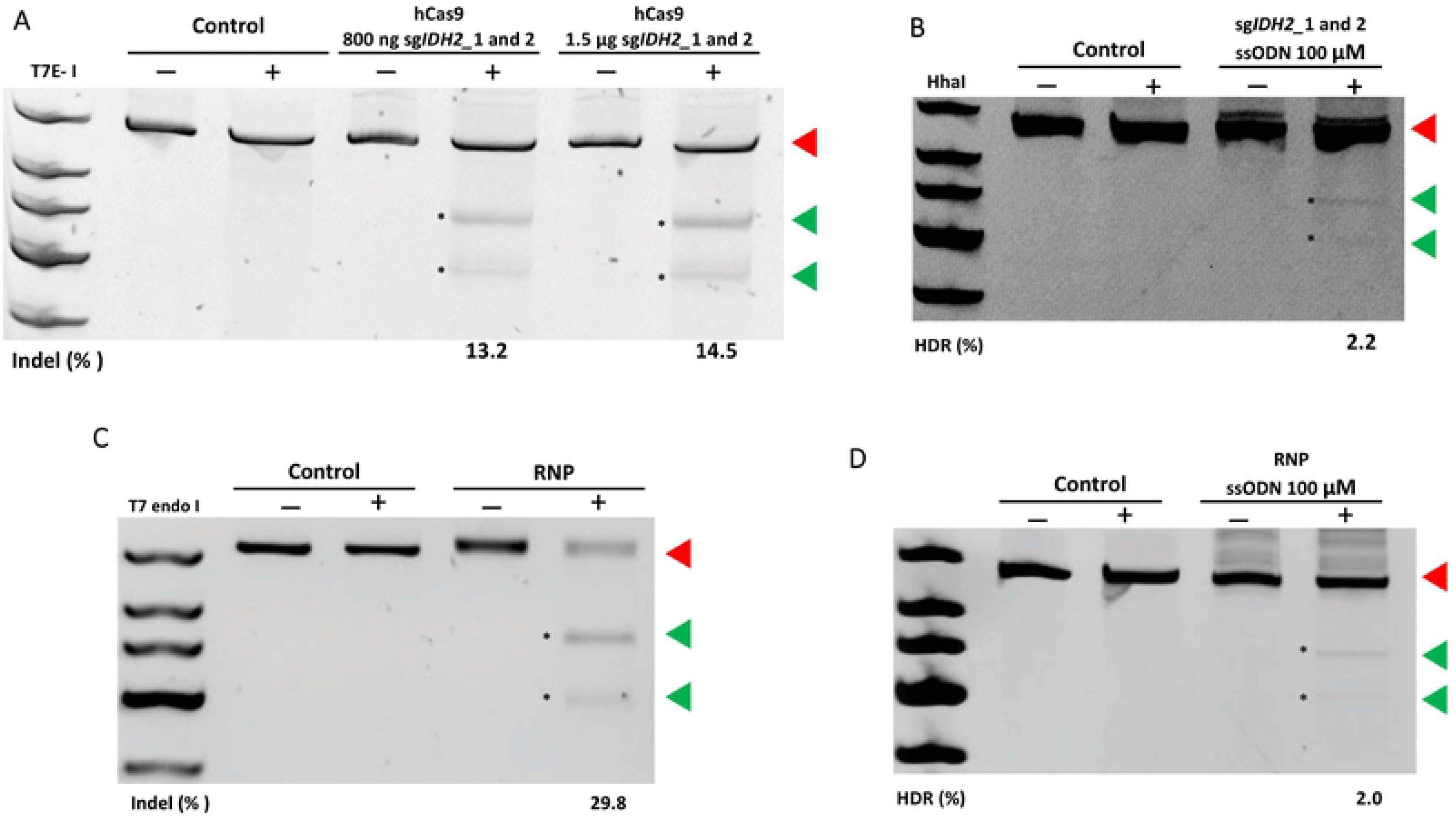
Analysis of *IDH2*^R172K^ mutation introduced by CRISPR into NB4 cells. **(A)** Electrophoresis in acrylamide gel of the products of the T7 endonuclease I assay, depicting indel formation by the sgRNA selected, using two different concentrations **(B)** Acrylamide gel to resolve the RFLP assay to investigate the rate of genome editing, using 100 μM of ssODN. The RFLP products were resolved in polyacrylamide gel. The cleaved products were used to quantify editing efficiency. For better visualization gel brightness and contrast were modified after quantification. **(C)** Acrylamide gel to resolve the products of the RFLP analysis, from cells targeted using RNP complexes. T7 endonuclease assay performed with RNA complexes with sg*IDH2*_1 and sg*IDH2*_2. **(D)** Acrylamide gel to resolve the RFLP assay to test edition in NB4 cells. Black numbers show band relative intensities (%). Red triangles point uncut products, and green triangles cut products. The percentage of cleavage with each combination of DNAs or RNPs is described below of each run (See full polyacrylamide gel in Figure S2 A-D).

Since our method to produce sgRNAs showed to be functional in NB4-Cas9 cells, we sought to compare the efficiency of these PCR-generated guides with RNP complexes. Transfection of RNPs complexes in NB4 cells produced 29.8% of indels (Figure 5C). With ssODN, we got a 2 % average of edition (Table 4) (Figure 5D), which is comparable to our method to produce CRISPR components.

### Deep sequencing of CRISPR-treated cells shows similar rates of gene editing and no off-targeting events

Amplicon deep sequencing was used to investigate gene editing efficiency and also potential off-target modifications, among the population of NB4-Cas9 and NB4 cells edited for *IDH2*^R172^ mutation. Sequencing data was analysed using the CRISPResso 2 software. The sequencing reads, are shown in Figure 6. For the NB4-Cas9 edited cells, 10.95 % of reads showed NHEJ repair, and 22.37 % for NB4 edited cells. The main modifications observed in the *IDH2 locus* were deletions. NB4-Cas9 edited cells displayed mainly deletions in the range of 20-29 pb (5.80 % of total reads). In the same way, NB4 edited cells showed mostly deletions of the same length (10.52 % of total reads), followed by deletion of 30-39 pb (10.20 % of total reads). The prevalence of reads with a precise DNA deletion between each PAM sequence is a characteristic result of combined use of two sgRNA [28].

**Figure 6.**
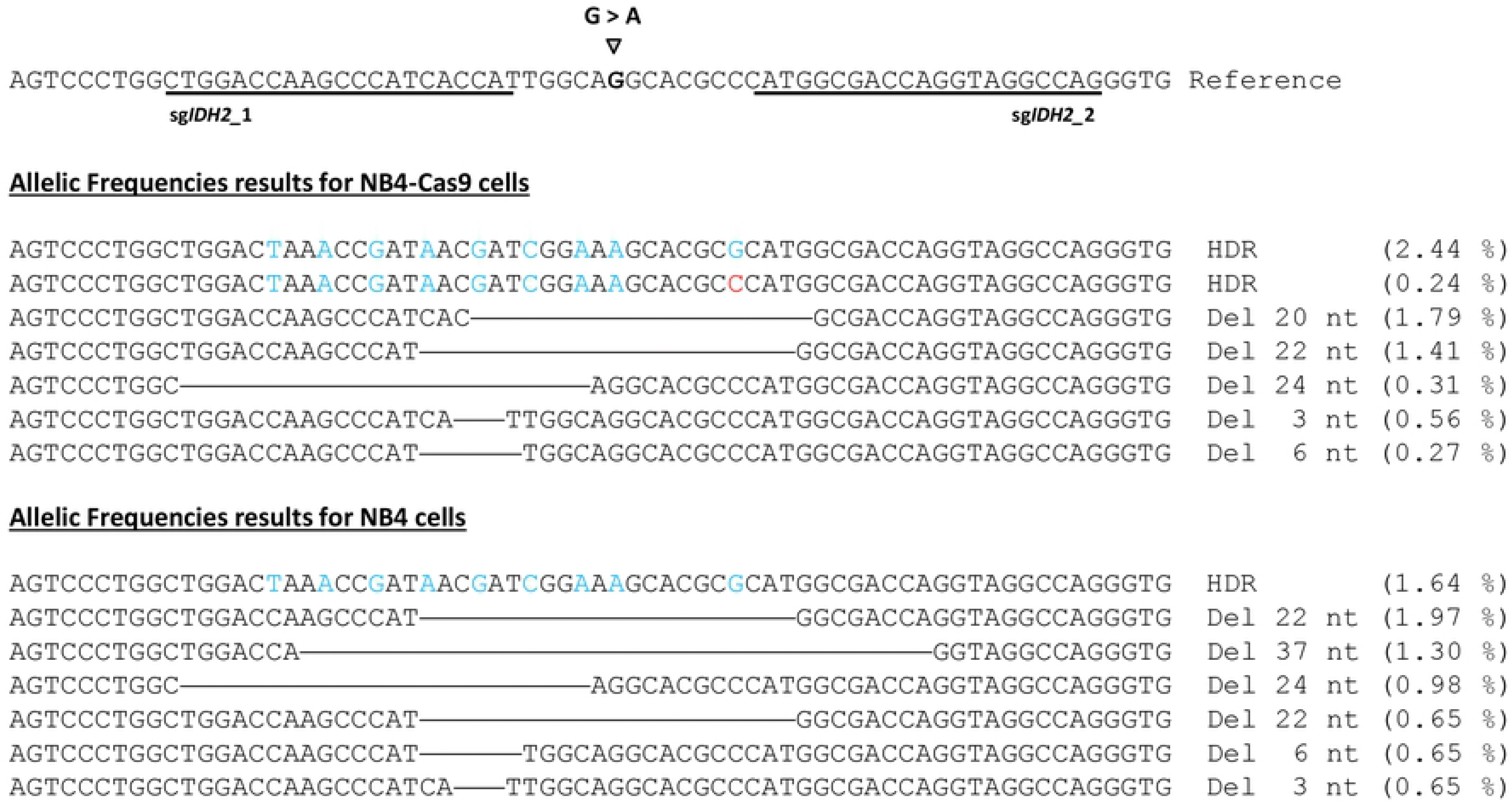
Summary adapted from CRISPResso2 results of the main identified alleles generated from edition *IDH2* gen. In the first line of each experiment is shown the reference sequence with the two sgRNA used underlined and the desire mutation to introduce. Deletions are shown in horizontal lines. Blue nucleotides are resulted from gene editing by ssODN. Red nucleotide represents an imperfect HDR.

In regard of knock-ins for the edited NB4-Cas9 cells, we obtained 2.44 % edited reads. Within that percentage, 0.42 % of reads integrated all ssODN changes less *IDH2*_2 PAM modification. These successful events of gene editing remove the HhaI site. In the case of edited naïve NB4 cells, we obtained 1.64 % of edited reads. Furthermore, CRISPResso2 classified reads with partial homology directed repair (HDR) in different editing subtypes. Reads with ssODN changes and other type of modifications (deletions or insertions) were classified as imperfect HDR (less than 0.5 % of reads). Finally, 0.19 % and 5.59 % of reads were filed as ambiguous reads for NB4-Cas9 and NB4 edited pools, respectively. These reads showed deletions with the extension of 22-99 pb, altering the quantification window used by the software. Due to this, these ambiguous reads are not classified as NHEJ repaired reads.

If we consider all reads with any kind of change as Cas9 activity, the results are comparable to the T7E-I assays. For NB4-Cas9 edited cells we obtained 14.52 % of indels and 13.82 % of edited reads by Next Generation Sequencing (NGS). For the NB4 edited cells, we observed 29.8 % of indel production, as shown by the T7E-I assay, and 29.9 % of edited reads by NGS. No significant percentages of reads with changes were detected in the potential off-targets in this mass sequencing analysis.

## DISCUSSION

In haematology, the CRISPR/Cas 9 system has triggered the development of many *in vitro* and *in vivo* models, along with new therapies (González-Romero et al). The main CRISPR vehicle used for *in vitro* experiments with haematopoietic cells are lentivirus. In the published literature, using haematological cells as models of leukaemia with lentivirus, we found cut efficiencies in range of 10 [14] to 90 % [11]. And up to 10 % for edition when using a DNA template for recombination [14]. To elude the major disadvantages of using lentivirus, we have developed a quick strategy to edit genes in leukemic cells.

We developed the pEGR1 vector and a protocol to produce sgRNAs by PCR. Using this, we produced sgRNA against *MYBL2* and *IDH2* and tested their functionality in HEK293 cells. Our PCR sgRNA, co-transfected with the hCas9 vector, showed a tendency to produce higher number of indels than standard CRISPR plasmids, like PX458, in HEK293. Therefore, our new method, which do not require bacterial transformation and selection, may be an alternative to using big CRISPR plasmids in these cells. On the other hand, this strongly suggests that reducing the size of the DNA increases substantially the chances of gene targeting. To edit *IDH2*, we combined two sgRNAs, which very much increased DSB than transfecting each guide individually, in agreement with other works [29].

Despite we obtained moderate efficiencies in HEK293, these results are in line with others published elsewhere [30–32], although many groups reported higher results in this cell line [28,33,34]. This discrepancy is probably due to many factors as cell type and target specific issues [31]. Many works proved that eukaryotic chromatin structures are critical factors for Cas9 nucleases efficiency [35–38]. Eukaryotic DNA forms complex structures as nucleosomes, which block Cas9 interaction with PAM sequences [35]. Isaac and colleagues reported than CRISPR/Cas9 system is less efficient for targeting *loci* located at entry/exit sites of nucleosomes and in nucleosomal dyad than in nucleosome flanking regions [35]. Nucleosome structures are not static and are cell type specific [39]. So, a difficult accessibility of *MYBL2* and *IDH2* DNA sequences in HEK293 could explain these modest efficiencies.

Finally, we replaced the hCas9 vector with PCR-amplified Cas9 cassette, although the indel production was reduced. In this regard, it has been described that lipofectamine forms compact structures with circular DNA, and this is what induces cell uptake. However, long linear DNA forms necklace-like structures that disrupt uptake by cells [40,41]. Furthermore, linear DNA is more vulnerable to exonuclease enzymes due to their unprotected ends [42]. Altogether, this explain low editing efficiency using these PCR-generated Cas9 cassettes.

When we used NB4 cells, we were barely able to introduce the PX458 vector. Hence, we decided to produce stable cell lines that constitutively expresses the Cas9 gene using a lentivirus, which encodes the nuclease. Three cell lines that express Cas9 were developed in our group (NB4-Cas9, HL60-Cas9 and MOLM-13-Cas9). Although we only used NB4-Cas9 in this work, these stable cell lines will allow us develop different stable models to study leukemic progression and the associated transcriptional changes. These lines were constituted with just one step of lentivirus transduction, since later steps of CRISPR transfections would include the constructs encoding the sgRNAs. Therefore, these future transfections do not include the uses of viral vectors, hence avoiding extra insertions of DNA. Altogether, this provides us with a model to produce CRISPR with easy, not only to do the proof-of-concept described here, but also for future investigations. Using flow cytometry, we reported higher transfection ratios with our GFP-containing guides, compared to PX458 (around 10 kb) or pMAX (around 3 kb) vectors, in the NB4-Cas9 cells. Our results are in agreement with the work of Wu and colleagues, who describe that PCR amplified EGFP produces high nucleofection efficiency in NB4 cells than the parental plasmid, with little effect in cell viability [43]. Targeting of *IDH2* produced 10.52 % of cuts and 2.2 % of editing efficiency using our technology. NB4 cells are really difficult to transfect. According to commercial nucleofection protocols, in ideal conditions, it is possible to obtain nearly 80 % of transfection efficiency using pMAX vector. But in our hands, we achieved 26 % with the same vector after many optimizations. Hence, we have reached the range of editing efficiencies for other leukemic cells, as the easy to transfect K562. This cell line is often used for gene editing optimization prior to edit stem cells due to its high transfection rate [11,33].

RNPs are usually chosen for gene editing due to their high efficiency. RNPs are recommended in cases where predicted potential off-targets, since they have a reduced window of activity. In our hands, RNPs increased indel occurrence but this did not correlated with higher editing rates, which agrees with previous reports [44]. This inconsistency may be caused by the cellular DNA reparation pathway, which tends to repair any genomic lesion, to avoid genomic instability [45]. HDR, using the sister chromatid as template, is the preferred way of fixing lesions, because it ensures reliable reparation [46], but NHEJ is easier and faster [47]. Hence, the molecular characteristics of both repair ways may contribute to a higher frequency of NHEJ correction.

Using NGS, we analysed 16 potential off-targets in edited NB4-Ca9 with our constructs and NB4 cells edited with RNPs. No off-target effects were detected, corroborating our idea that Cas9 gene inserted in NB4-Cas9 genome is secure.

## CONCLUSION

In this work we have expanded the CRISPR/Cas9 toolkit by developing a robust and alternative strategy to generate CRISPR constructs. Even though we have focused in NB4 cells, this technology could be used to edit diverse classes of cell lines.

## ACKNOWLEGMENTS

We would like to thank Lourdes Cordón, from the Citomic Unit of the HULAFE.

## Funding

An Integrated Project of Excellence, from the Instituto de Salud Carlos III (ISCIII, Madrid, Spain), mostly funded this work (PIE13/00046). RPVM is a Miguel Servet type II researcher (CPII16/00004) ISCIII. This study was supported in part by research funding from FEDER (CIBERONC, CB16/12/00284). Grants from the “Instituto de Salud Carlos III” (ISCIII) fund the laboratory of RVM and JVC (PI14/00949, PI17/00011, PI16/01113, PI/17/0575, and PI19/00812). All grants from ISCIII are cofinanced by the European Development Regional Fund ‘’A way to achieve Europe’’ (ERDF). Some equipment used in this work have been funded in partnership between the Generalitat Valenciana (Conselleria de Sanitat I Salut Pública, Valencian Community, Spain) and European Funds (ERDF). The funds from the ISCIII are partially supported by the European Regional Development Fund. RPVM is also a Marie Curie fellow (CIG322034, EU). EGR is a recipient of a FEHH 2020 fellowship.

## Author’s contributions

Conceptualization, E.G.-R., R.P.V.-M. and J.V.C.; Methodology, E.G.-R, R.P.V.-M., A.L. and G.G.-G.; Investigation, E.G.-R., A.R.-V., C.M.-V. and A.L.; Writing – Original Draft, E.G.-R; Writing – Review and Editing, R.P.V.-M., A.L., G.G.-G., J.V.C., J.M.M. and M.A.S. All authors have collaborated in the production of the manuscript, and agree with its publication.

## SUPPORTING INFORMATION

**S1.** sgRNAs, ssODN and primers sequences.

## Notes

### Competing Interest Statement

The authors have declared no competing interest.

